# *In-vitro* pathogen inhibition: Comparing the inhibitory effects of a complex multistrain synbiotic with simple probiotics containing the yeast *Saccharomyces boulardii or Lactobacillus rhamnosus bacteria*

**DOI:** 10.1101/543819

**Authors:** Jacek Piatek, Henning Sommermeyer, Arleta Ciechelska-Rybarczyk, Malgorzata Bernatek

**Author notes:** Corresponding author (JP). These authors contributed equally to this work.

## Abstract

Supplementation with probiotics is considered as alternative treatment or adjuvant therapy for a number of bacterial infections for which the use of antibiotics is either not recommended or emerging antibiotic resistance is a major concern. Inhibition of the growth of pathogenic bacteria has been related to a number of different activities of probiotic bacteria or yeasts, some of which are very specific for particular strains of probiotics. As the different inhibition activities might act additively or even synergistically, probiotic multistrain products are discussed as potentially being more effective in pathogen inhibition than products containing one or a small number of probiotic strains. The present study investigated the in vitro inhibition of *Escherichia (E.) coli, Shigella spp.*, Salmonella (*S.) typhimurium* and *Clostridum (Cl.) difficile*, all being human pathogens of significant worldwide healthcare concerns. The probiotic containing the yeast *Sacharomyces (S.) boulardii* inhibited all four pathogens. Similar inhibitions were observed with a bacterial probiotic containing three different strains (Pen, E/N and Oxy) of *Lactobacillus (Lc.) rhamnosus.* Compared to the inhibition found for these probiotics, the inhibitory effects of a complex multistrain synbiotic, containing nine different probiotic strains (6 *Lactobacilli* and 3 *Bifidobacteria*) and the prebiotic *fructooligosaccharide (FOS),* were significantly stronger. The stronger inhibition by the complex multistrain synbiotic was observed for all four tested pathogens. Our findings support a hypothesis that complex synbiotic products containing a larger number of different strains combined with a prebiotic component might be more attractive candidates for further clinical characterization than simpler probiotics containing one or only few probiotic strains.

## Introduction

While antibiotic therapy appropriately prescribed is highly effective in eliminating infections by bacterial pathogens, the therapy also alters the microbial balance in the gastrointestinal tract. This disturbance of the gut microbiota can cause symptoms (e.g., antibiotic-associated diarrhea) but is frequently not noticed by the patient. Even without symptoms, the altered gut microbiota might cause long term effects, e.g. an altered function of the patient’s immune system. About 75% of the human immune system is closely associated with the gastrointestinal tract and the gut microbiota is vital for its development and function. A diverse and well balanced gut microbiota is essential to keep the growth of potentially pathogenic germs in the inner part of the gastrointestinal system under control. An unbalanced and/or diversity-reduced gut microbiota can result in proliferation of pathogenic germs, overgrowth of the gut by the pathogens and finally lead to manifestation of diseases.

Colonization by potentially pathogenic germs, due to infection with these bacteria or the use of antibiotics, can be limited or prevented by the administration of probiotic bacteria. The probiotic germs limit the proliferation of pathogenic bacteria by a variety of mechanisms. Among these is secretion of organic acids, *bacteriocins, bacteriocin*-like substances, as well as the inhibition of the adhesion of pathogenic bacteria to the gut epithelium. Probiotics are available in the form of yeast probiotics containing e.g. *S. boulardii* or bacterial probiotics. Bacterial probiotics can be further differentiated into simple probiotics (containing only one or a small number of probiotic bacteria) or complex multistrain probiotics (containing a larger number of different probiotic bacteria). Synbiotics contain, in addition to the probiotic component(s), a prebiotic ingredient, e.g., *fructooligosaccaride*, which can be used by probiotic germs as a source of energy.

Probiotics are discussed as alternative treatments or adjuvant therapy for a number of bacterial infections for which use of antibiotics is either not recommended or emerging antibiotic resistance is of concern. Monostrain probiotics containing the yeast *S. boulardii* [1–3] or the bacterium *Lc. rhamnosus* [4–6] have been extensively characterized in the scientific literature. While monostrain probiotics are commonly used in medical practice, there is a recent trend to shift to complex multistrain probiotics [7,8] and/or complex multistrain synbiotics [9]. The underlying driver for this shift is the assumption that more strains are enhancing the chance of therapeutic success, as they are providing a broader spectrum of pathogen inhibitory effects. In addition, a mixture of probiotic bacteria might express synergistic effects, which exceed the sum of the contributions of each individual probiotic bacterium.

The present study compared the in-vitro pathogen inhibition of a yeast monostrain probiotic *S. boulardii*, a bacterial probiotic containing three (Pen, E/N, Oxy) strains of *Lc. rhamnosus* and a complex multistrain synbiotic product containing 9 different probiotic bacteria (6 *Lactobacilli* and 3 *Bifidobacteri*a) and a prebiotic compound (*fructooligosaccharide*). Inhibitory effects of the three products were evaluated against *E. coli, Shigella spp., S. typhimurium* and *Cl. difficile*.

## Materials and methods

### Products tested

#### *S. boulardii* probiotic

The yeast monostrain *S. boulardii* probiotic product used in the experiments is commercially available as Enterol 250 (capsules each with 250 mg of freeze-dried S. boulardii) and was obtained from Biocodex, 7 avenue Gallieni, 94250 Gentilly, France. The product contains the S. boulardii strain CNCM I-745.

#### *Lc. rhamnosus* probiotic

The bacterial *Lc. rhamnosus* probiotic product is commercially available as Lakcid (sachets each with a total of 2 x 10^9^ cfu of bacteria) and was obtained from Biomed-Lublin, ul. University 10, 20-029 Lublin, Poland. The product contains a mixture of the Lc. rhamnosus strains Pen (40%), E/N strain (40%) and Oxy (20%) [10].

#### Complex multistrain synbiotic

The complex multistrain synbiotic product is commercially available as Multilac^®^ Baby (stickpack with a total of 1 x 10^9^ cfu of bacteria) and was obtained from Vivatrex GmbH, Martinstr. 10-12, 52062 Aachen, Germany. Each stickpack of the product contains: 1.11 x 10^8^ cfu *Lactobacillus acidophilus LA-14* [11, 12]; 1.11 x 10^8^ cfu *Lactobacillus casei R0215*; 1.11 x 10^8^ cfu *Lactobacillus paracasei Lpc-3*; 1.11 x 10^8^ cfu *Lactobacillus plantarum Lp-115* [11, 13–21]; 1.11 x 10^8^ cfu *Lactobacillus rhamnosus* [4–6], 1.11 x 10^8^ cfu *Lactobacillus salivarius Ls-33*, 1.11 x 10^8^ cfu *Bifidobacterium lactis Bl-04* [22], 1.11 x 10^8^ cfu *Bifidobacterium bifidum R0071* [23,24], 1.11 x 10^8^ cfu *Bifidobacterium longum R0175* [25,26]. The prebiotic ingredient of Multilac^®^ Baby is fructooligosaccharide (1.43 g/stickpack).

### In-vitro pathogen inhibition experiments

#### Inhibition of *E. coli, Shigella spp*. and *S. typhimurium*

For the in-vitro pathogen inhibition experiments with *E. coli* (strain 25922), *Shigella spp.* and *S. typhimurium* (strain 14028), the respective pathogen was inoculated on Columbia agar with 5% sheep blood and incubated at 37° C under aerobic conditions for 24 h. Suspensions of the three products (*S. boulardii* probiotic, *Lc. rhamnosus* probiotic and the multistrain synbiotic) were inoculated with a density of 2 on the McFarland scale on MRS medium (Oxoid) and incubated for 48 h in the presence of 5% CO2. After incubation in the MRS medium, 10 mm diameter bars were cut out and transferred to a Mueller-Hinton medium (Oxoid) previously inoculated with the respective pathogen strain with a density of 2 on the McFarland scale. The tested cultures were stored at 4° C for 4 h, followed by an incubation at 37° C for 24 h under aerobic conditions.

#### Inhibition of *Cl. difficile*

For the in-vitro pathogen inhibition studies with *Cl. difficile*, the pathogen was cultivated under anaerobic conditions at 35-37° C for 24-48 h on Schaedler medium (Oxoid). Suspensions of the three evaluated products were inoculated with a density of 2 on the McFarland scale on MRS medium (Oxoid) and incubated for 48 h in the presence of 5% CO2. 10 mm diameter bars were transferred to a Mueller-Hinton medium (Oxoid) with horse blood and NAD (Oxoid) and incubated under anaerobic conditions for 24 h.

### Measurement of growth antagonism

Measurement of growth antagonisms between the different pathogenic bacteria and the evaluated two probiotics and the complex multistrain synbiotic were calculated by means of the bar graph method according to Strus [27,28]. At the end of the incubation, measurements of inhibition zones around the tested colonies were taken from the outer edge of the colonies to the outer edge of the clear zones. Each test was performed in triplicate and the arithmetic means of the radii measuring from the edges of the colonies to the edges of the clear zones were calculated as well as the standard deviations SD. Statistical differences between the inhibition of the three different products was evaluated by using the independent group T-test (significance p<0.01). Tables containing all raw data from the in-vitro pathogen inhibition experiments, the arithmetic means and standard deviations calculated are provided in S1 Dataset.

## Results and discussion

### Inhibition of *E. coli*

*E. coli* was inhibited by all three tested products. Inhibition by *S. boulardii* was 6.0±1.0 mm, by *Lc. rhamnosus* 3.7±0.6 mm, and by the complex multistrain synbiotic 13.7±1.2 mm (Fig 1). There was no significant difference between the inhibition by *S. boulardii* and *Lc. rhamnosus* (p=0.0125). Inhibition by the complex multistrain symbiotic was significant stronger than the inhibition by the probiotics. P-values compared to *S. boulardii* and to *Lc. rhamnosus* were p=0.0005 and p=0.0001, respectively.

**Fig 1.**
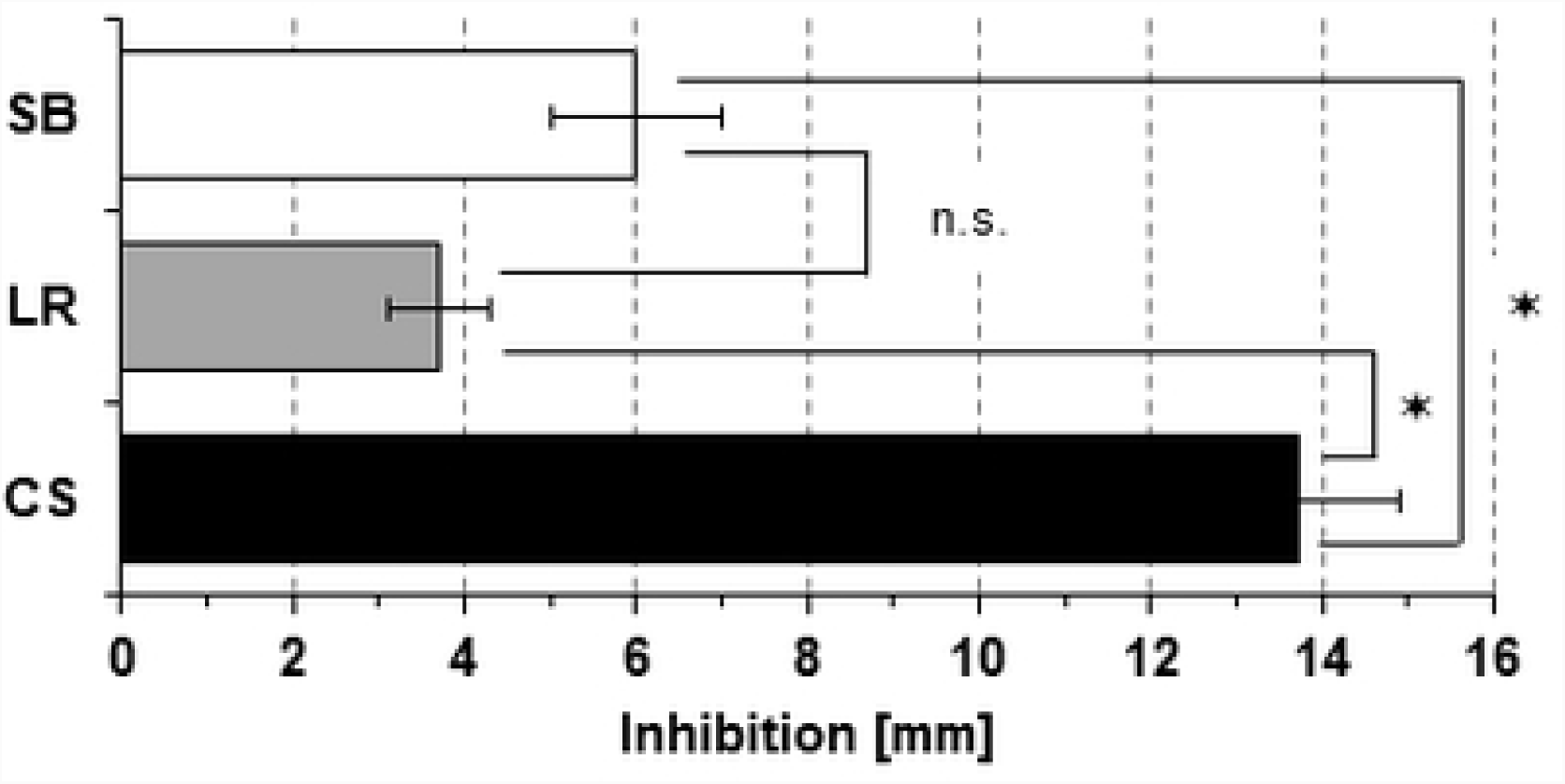
In-vitro inhibition of *E. coli.* Inhibition by yeast S. boulardii probiotic (SB), Lc. rhamnosus probiotic (LR) and a complex multistrain synbiotic (CS). Bars represent the arithmetic mean of three independent experiments, error bars the standard deviation SD. ⋆ indicates a significant difference (p-value <0.01) while n.s. indicates no significant difference (p-value>0.01).

While most of *E. coli* isolates from the human intestine are essential and beneficial for the human body, there are also pathogenic *E. coli* species. Therefore, *E. coli* has to be considered as an important cause of diarrheal illness, with a variety of phenotypes and pathogenic mechanisms. As long as there is no evidence for a systemic infection, antibiotic therapy is rarely indicated and should be deferred until culture results are available. Due to emerging resistance of *E.coli* against antibiotics [22,29,30], probiotics are discussed as alternative treatments or adjuvant therapy against *E. coli* infections. *S. boulardii* has been found to have a number of positive effects in different experimental *E. coli* pathogen models [1, 31–38]. In vitro pathogen inhibition of E. coli by S. boulardii has been investigated [39] but no clear antagonism could be demonstrated. Lc. rhamnosus has been demonstrated to inhibit growth of E. coli in vitro [40]. Similar effects have been found for a number of monostrain probiotics containing either *Lactobacilli* or *Bifidobacteria* [7,22,40–45]. In a recent study comparing the in vitro antagonistic activity of monostrain and multistrain bacterial probiotics against *E. coli*, it was found that multistrain probiotics exhibited a significantly stronger inhibitory effect than monostrain probiotics [22]. Results from the present study (Fig 1) supports the latter finding, as it was also found that a complex mixture of probiotic bacteria (combined with FOS) has a significant stronger inhibitory effect than less complex yeast or bacterial probiotics.

### Inhibition of Shigella spp.

All three tested products inhibited *Shigella spp.*. Inhibition caused by *S. boulardii* and *Lc. rhamnosus* were similar, 5.7±1.5 mm and 5.3±0.6 mm, respectively. The inhibition caused by the complex multistrain synbiotic was 13.7±1.5 mm (Fig 2). While there was no significant difference between *S. boulardii* and *Lc. rhamnosus* inhibitions (p=0.187), the effect of the complex multistrain symbiotic was significant stronger when compared with *S. boulardii* (p=0.0001) or *Lc. rhamnosus* (p=0.0005).

**Fig 2.**
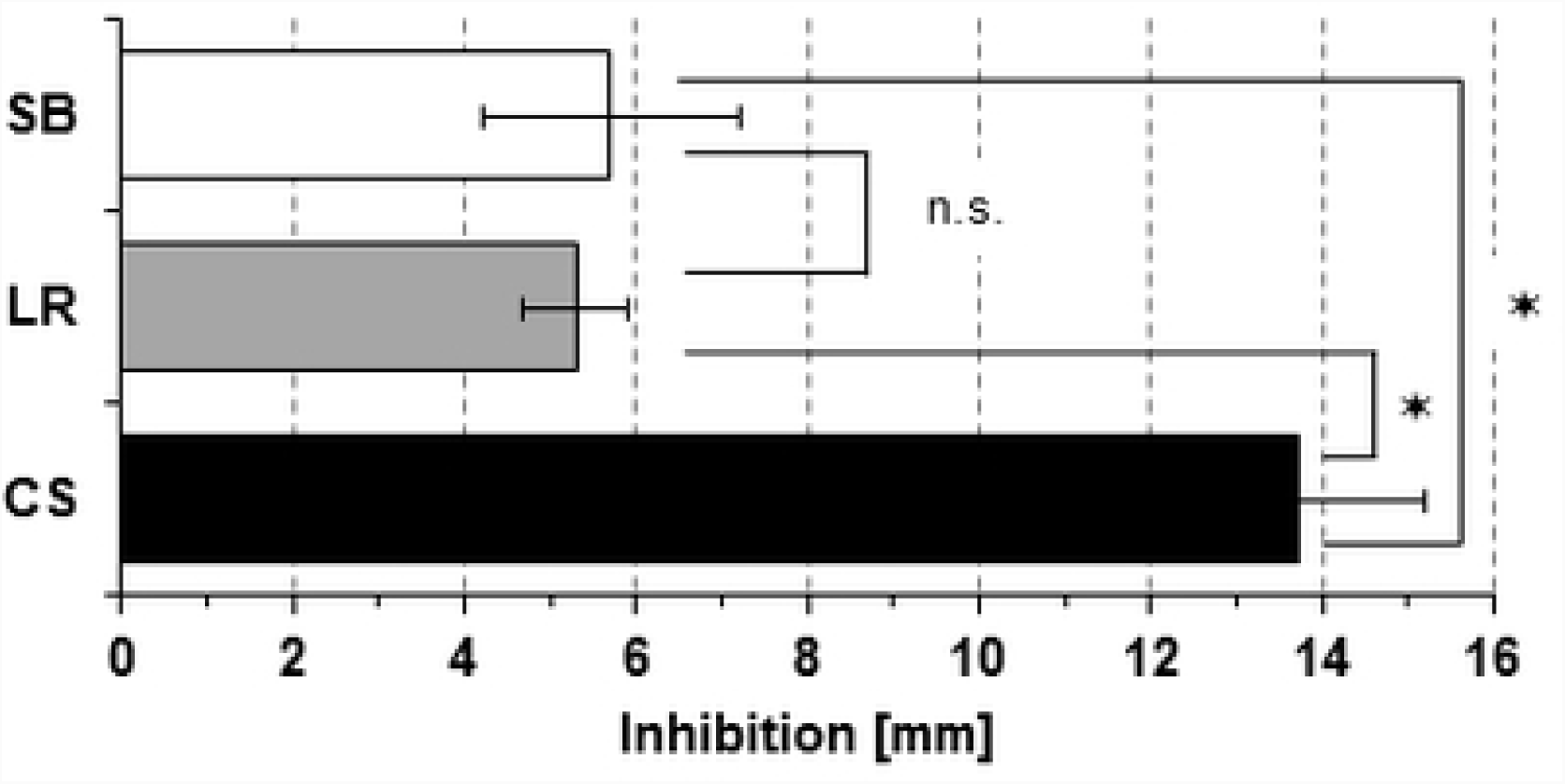
Inhibition of *Shigella spp.* Inhibition by yeast S. boulardii probiotic (SB), Lc. rhamnosus probiotic (LR) and a complex multistrain synbiotic (CS). Bars represent the arithmetic mean of three independent experiments, error bars the standard deviation SD. ⋆ indicates a significant difference (p-value <0.01) while n.s. indicates no significant difference (p-value>0.01).

*Shigella* infections are a major public health problem in areas of poor sanitation. *Shigellosis* is spread by fecal-oral transmission and ingestion of a small number of *Shigella* bacteria can already cause clinical disease. The pathology of a *Shigella* infection is characterized by an acute inflammation with polymorphonuclear cell (PMN) infiltration, resulting in massive destruction of the colonial mucosa and characteristic dysentery syndrome. The highest incidence is observed in children (< 5 years). Most patients recover from *Shigellosis* without treatment within 5-7 days. Various antimicrobial agents are effective in the treatment of severe cases of *Shigellosis*, however a globally emerging drug-resistance is observed [46]. Beneficial effects during *Shigella* infections have been described for *S. boulardii* [32,37,47,48], bacterial single-strain probiotics [7,40,49] and multistrain probiotics [7]. The present study found that *S. boulardii* and *Lc. rhamnosus* exhibit an inhibitory effect on the in vitro growth of *Shigella spp..* While the effect of both, the yeast monostrain and the Lc. rhamnosus probiotic, were small and similar in extent, the tested complex multistrain synbiotic product resulted in a significantly stronger growth inhibition.

### Inhibition of S. typhimurium

Inhibitory effects of the three products in individual inhibition experiments with *S. typhimurium* exhibited a higher variability compared to the inhibition of the other three tested pathogens. Inhibition by *S. boulardii* was 5.0±1.7 mm, by *Lc. rhamnosus* 5.0±2.0 mm and by the complex multistrain synbiotic 12.0±2.0 mm (Fig 3). No significant difference between the inhibition by *S. boulardii* and *Lc. rhamnosus* was found (p=0.5). Inhibition by the complex multistrain symbiotic was significant stronger when compared with *S. boulardii* (p=0.0050) or *Lc. rhamnosus* (p=0.0065).

**Fig 3.**
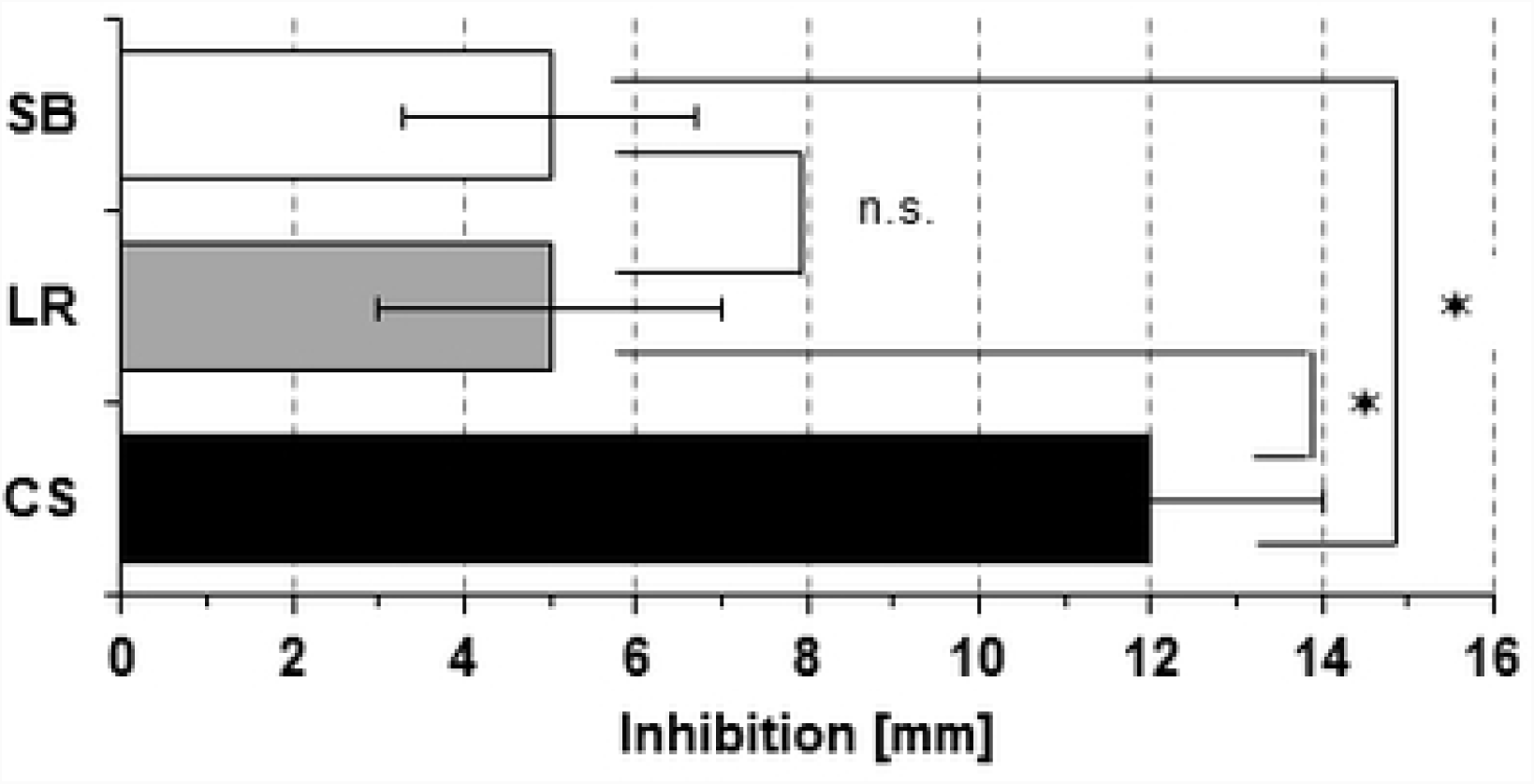
Inhibition of *S. typhimurium.* Inhibition by yeast S. boulardii probiotic (SB), Lc. rhamnosus probiotic (LR) and a complex multistrain synbiotic (CS). Bars represent the arithmetic mean of three independent experiments, error bars the standard deviation SD. ⋆ indicates a significant difference (p-value <0.01) while n.s. indicates no significant difference (p-value>0.01).

Most food-borne bacterial gastroenteritis is caused by ingestion of food or water contaminated with *non-typhoidal Salmonella* (NTS). In most cases, NTS is a self-limiting disease that causes mild gastroenteritis. However, there are also cases of persistent NTS, in which the infection and its symptoms manifest for years [50]. The pathology of *Salmonella* infections is characterized by diarrhea often with heme-positive stool and abdominal pain. Excess antibiotic use in years prior to the infection is associated with a higher incidence of NTS. Disruption of the balance of the gut microbiota (e.g., by antibiotics) is discussed as a possible cause. For most of patients an antimicrobial therapy is not indicated. Positive effects on *Salmonella* infections have been described for *S. boulardii* [32,39,51,52], bacterial monostrain probiotics [7,40,41,43,53,54] and multistrain probiotics [7,55]. The present study confirms that *S. boulardii* and *Lc. rhamnosus* exhibit growth antagonistic properties against *S. typhimurium.* However, when the inhibitory effect of the probiotics was compared with that of a complex multistrain synbiotic, it was found that the inhibition by the complex multistrain synbiotic was significantly stronger.

### Inhibition of Cl. difficile

*Cl. difficile* was sensitive to inhibition by all three tested products. *S. boulardii* and *Lc. rhamnosus* had similar inhibitory effects (p=0.095), with inhibitions of 5.7±1.5 mm and 4.0±1.0 mm, respectively (Fig 4). The effect of the complex multistrain probiotic was significantly stronger when compared to *S. boulardii* (p=0.0001) or *Lc. rhamnosus* (p=0.0004).

**Fig 4.**
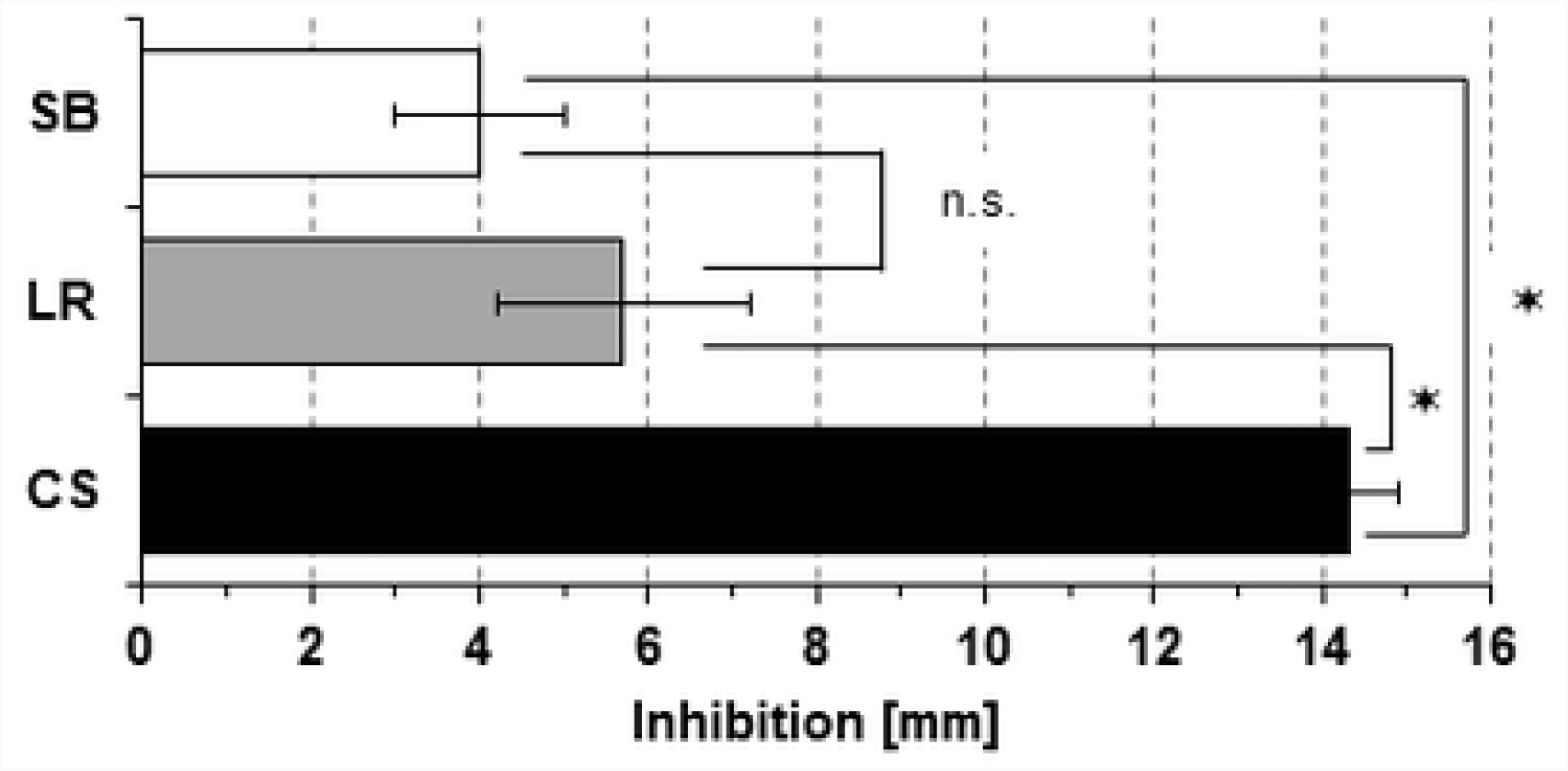
Inhibition of *Cl. difficile.* Inhibition by yeast S. boulardii probiotic (SB), Lc. rhamnosus probiotic (LR) and a complex multistrain synbiotic (CS). Bars represent the arithmetic mean of three independent experiments, error bars the standard deviation SD. ⋆ indicates a significant difference (p-value <0.01) while n.s. indicates no significant difference (p-value>0.01).

Exposure to antibiotics is the most important risk factor for an initial infection with *Cl. difficile* (CDI). Other risk factors are older age, hospitalization and gut disease preconditions of the patient. Certain antibiotics (e.g., clindamycin, quinolones, cephalosporines) are associated with a higher risk of causing CDI. Pathology of CDI is characterized by an antibiotic-caused disruption of the gut microbiota, resulting in an overgrowth by *Cl. difficile*, production of toxins and disease development. Characteristic symptoms are diarrhea and abdominal pain. In severe cases, the CDI can result in a life-threatening *pseudomembranous colitis*. Recurrence of CDI is a not uncommon observation. First step in the treatment of CDI is discontinuing the therapy with the inciting antibiotic as soon as possible. Depending on severity of the CDI, three antibiotics can be considered for therapy: *metronidazole, vancomycin* and *fidaxomicin* [56]. Smaller clinical trials have found that *S. boulardii* in combination with *vancomycin* reduced CDI in patients with recurrent CDI [57,58], while it had no effect on the recurrence in patients with primary CDI [57]. There is less evidence that *S. boulardii* alone prevent CDI [59,60]. A meta-analysis concluded that there was moderate efficacy for probiotics in prevention of CDI, but this analysis combined studies of many different probiotics or mixtures of probiotics in varied clinical settings [61]. No efficacy in the prevention of CDI was found in a larger randomized control trial using a four-strain probiotic (*Lactobacillus acidophilus CUL60, Lactobacillus acidophilus CUL 21, Bifidobacterium bifidum CUL20, Bifidum lactis CUL34*) in elderly inpatients [62]. Data are still insufficient to conclude what benefit which type of probiotic can have in what kind of CDI patients [63]. The present study finds in vitro inhibition of *Cl. difficile* by *S. boulardii* and *Lc. rhamnosus*. Compared with the inhibitory effect of the tested complex multistrain synbiotic the inhibitory effect of the probiotics was significantly weaker.

## Conclusion

The beneficial role of probiotics in the four pathogen infection situations described above remains still not fully understood. However, there is emerging evidence that probiotics might be of benefit as alternative treatments or adjuvant therapies. Both tested probiotics exhibited growth inhibiting properties against all four examined pathogens. No difference was found between the size of the inhibitory effect of the yeast monostrain probiotic and that of the bacterial Lc. rhamnosus probiotic. Compared to the probiotics, the complex multistrain synbiotic was found to result in significant stronger in vitro inhibition of the tested pathogens.

More clinical research will be needed to characterize the potential benefits of probiotic or synbiotic products in case of infections with *E. coli, Shigella spp., S. typhimurium* or *Cl. difficile*. However, results from the present study suggest that testing the effect of complex multistrain synbiotics might be particular interesting to investigate in clinical settings.

## Supporting information

**S1 Dataset. Raw data and calculated values.** Tables containing all raw data from the in-vitro pathogen inhibition experiments (E. coli, Shigella spp., Salmonella typhimurium, Clostridium difficile), the arithmetic means and standard deviations calculated.

**S1 Text. Abstract in German Language.**

